# Enhanced glymphatic CSF tracer influx during α2-adrenergic agonist anesthesia is independent of tracer injection duration

**DOI:** 10.64898/2026.05.29.728816

**Authors:** Tianyi Tong, Evan Newbold, Jiayi Wang, Emma Waight, Ashley Caudell, Antonio Ladrón-de-Guevara, Michael J. Giannetto, Lauren M. Hablitz, Maiken Nedergaard

## Abstract

The glymphatic system mediates brain-wide cerebrospinal fluid (CSF) transport and is highly sensitive to brain state. Experimental studies show that α2-adrenergic agonist– based anesthesia enhances glymphatic CSF influx, whereas isoflurane markedly suppresses it. However, it has been suggested that the reduced tracer influx observed during isoflurane anesthesia may reflect rapid clearance of tracer from the basal cisterns rather than genuine inhibition of glymphatic transport. To address this question, we compared conventional short-duration cisterna magna tracer injections with prolonged low-rate infusion while maintaining identical total tracer dose and anesthesia duration. Across both paradigms, ketamine/dexmedetomidine anesthesia consistently produced substantially greater perivascular CSF influx than isoflurane. In contrast, tracer accumulation in blood and cervical lymph nodes remained largely unchanged between conditions. These findings demonstrate that suppression of glymphatic influx during isoflurane anesthesia is independent of tracer injection duration and support the conclusion that α2-adrenergic agonist–based anesthesia promotes glymphatic transport through mechanisms linked to sleep-like brain states.

## Introduction

The glymphatic system is a brain-wide fluid transport mechanism whose activity is strongly regulated by the sleep–wake cycle(1). Sleep enhances brain fluid transport, whereas wakefulness suppresses it (2, 3). Many preclinical studies of glymphatic function rely on cerebrospinal fluid (CSF) tracer injections into the cisterna magna. Because this approach is invasive, anesthetic regimens have commonly been used as surrogates for wakefulness and sleep. Isoflurane has served as a surrogate for wakefulness because it produces a low-amplitude, high-frequency EEG pattern, whereas α2-adrenergic agonists such as dexmedetomidine or xylazine, combined with ketamine, have been used as surrogates for sleep due to their high-amplitude slow-wave EEG activity (4-7).

These studies have consistently shown that isoflurane is associated with reduced CSF tracer influx into the brain, whereas dexmedetomidine/ketamine or xylazine/ketamine markedly increase tracer influx. However, it has been argued that isoflurane promotes rapid washout of CSF tracers from the basal cisterns, thereby artifactually reducing apparent brain influx. To directly test this possibility, we compared here the traditional 5-minute CSF tracer injection paradigm with a continuous 40-minute infusion protocol. Our analysis demonstrated that isoflurane suppresses brain CSF tracer influx independently of injection duration or paradigm compared with dexmedetomidine/ketamine. These findings support the conclusion that anesthetic regimens producing a sleep-like EEG pattern, particularly α2-adrenergic agonist-based anesthesia (5), enhance glymphatic CSF influx compared with isoflurane anesthesia.

## Results

We tested whether the fluorescent tracer injection paradigm into the cisterna magna influences the well-established difference in CSF influx between isoflurane- and ketamine/dexmedetomidine (K/Dex)-anesthetized mice (4-7). Across all groups, the total tracer volume, tracer concentration, and overall anesthetic duration were kept identical (Fig. 1A). Overall, the standard injection paradigm (2µl/min for 5 minutes) produced greater CSF influx into the perivascular spaces surrounding the middle cerebral artery, visualized in dorsal whole-brain images, compared with the prolonged low-rate infusion paradigm (0.25µl/min for 40 minutes). Importantly, K/Dex-anesthetized mice consistently exhibited greater CSF influx than isoflurane-anesthetized mice under both injection conditions (Fig. 1B). These findings indicate that K/Dex anesthesia is more permissive to perivascular CSF influx independently of injection duration or infusion rate. Similar differences were observed in the ventral whole brain (Fig. 1C) and across six coronal brain sections (Fig. 2).

**Figure 1:**
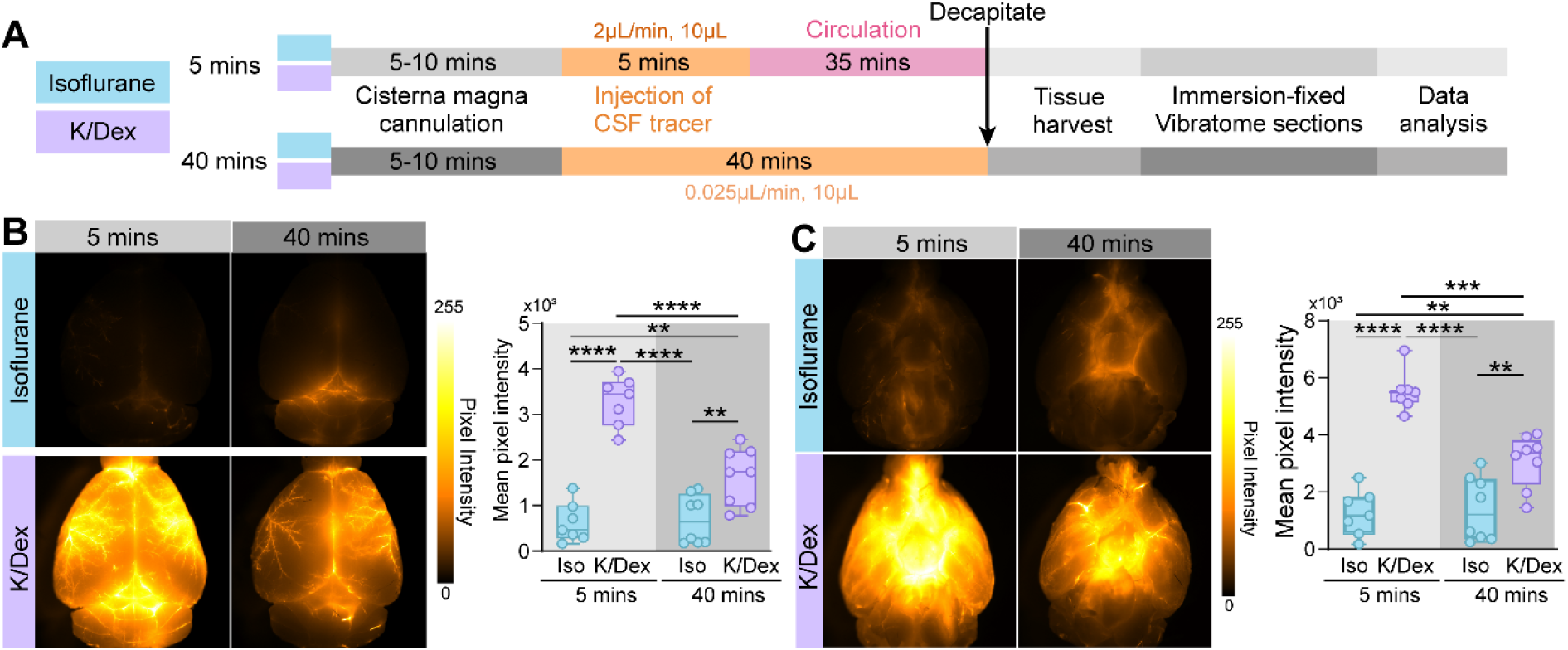
Ketamine/dexmedetomidine anesthesia enhances perivascular influx across the brain surface. (A) Schematic of experimental design. (B) *Left:* Representative dorsal whole-brain images obtained following either isoflurane or ketamine/dexmedetomidine (K/Dex) using either a 5 minute (5 mins) or 40 minute (40 mins) intracisternal tracer infusion. *Right*: min/max boxplots showing mean fluorescence intensity across the dorsal brain. (C) same analysis as in (B) for ventral whole-brain images. For all boxplots: the lower and upper whiskers indicate minimum and maximum values; the center line represents the median and individual circles represent individual mice. All statistics two-way ANOVA with Tukey HSD post-hoc tests. \*\**p*<0.01,\*\*\**p*<0.001,\*\*\*\**p*<0.0001. n = 7-8 mice per group. Complete statistical results are provided in the accompanying table.

**Figure 2:**
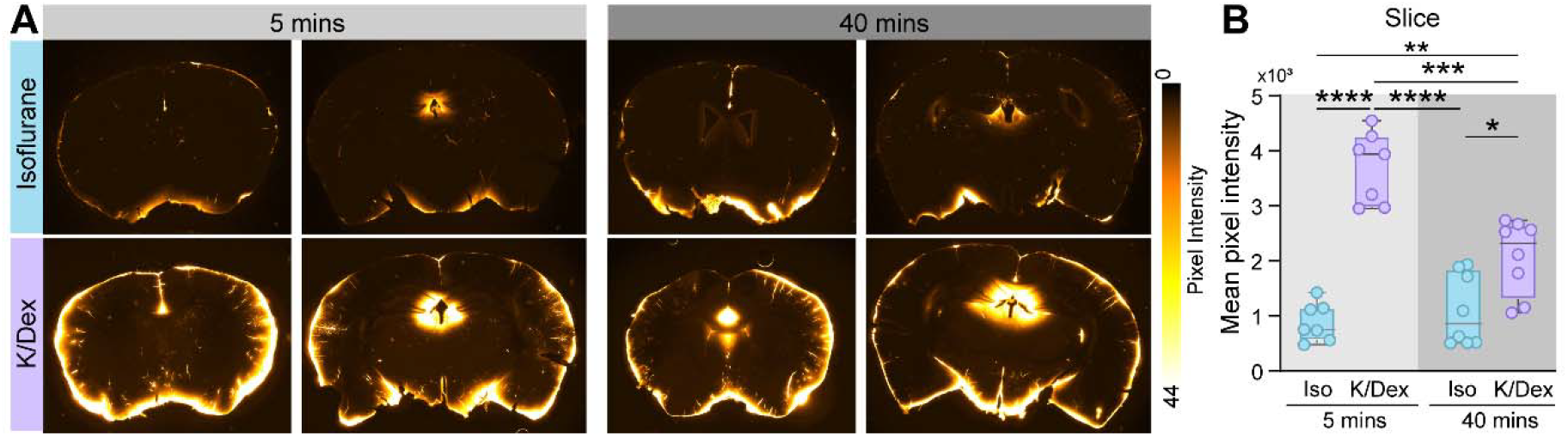
Ketamine/dexmedetomidine anesthesia enhances perivascular influx in coronal brain sections. (A) Representative coronal brain sections obtained from mice anesthetized with either isoflurane or ketamine/dexmedetomidine (K/Dex) after a 5 minute (5 mins) or 40 minute (40 mins) intracisternal tracer infusion. (B) min/max boxplots showing the averaged mean fluorescence intensity across coronal sections. For all boxplots: the lower and upper whiskers indicate minimum and maximum values; the center line represents the median and individual circles represent individual mice. Statistical analysis was performed using two-way ANOVA with tukey HSD post-hoc tests. \**p*<0.05, \*\**p*<0.01, \*\*\**p*<0.001, \*\*\*\**p*<0.0001. n = 7-8 mice per group. Complete statistical results are provided in the accompanying table.

There is a growing interest in efflux pathways by which CSF exits the cranial cavity. To assess whether anesthetic or tracer injection paradigm influence CSF efflux routes, dorsal skull, blood serum, and both superficial and deep cervical lymph nodes (scLN and dcLN, respectively) were collected under the same anesthetic and injection paradigms described in Fig. 1A. In the dorsal skull with the dura intact (Fig. 3A, Fig. 3B top), K/Dex anesthetized mice exhibited greater tracer accumulation than isoflurane-anesthetized mice, without significant effects of injection paradigm. In contrast, no significant differences were detected in blood serum fluorescence intensity after 40 min of tracer circulation (Fig. 3B bottom), or in average fluorescence signal within the lymph nodes (Fig. 3CD). A non-significant trend (*p* = 0.0696) towards greater tracer accumulation in the deep cervical lymph nodes with K/Dex was observed following the 5-minute injection paradigm (Fig. 3D bottom), potentially consistent with enhanced perivascular CSF influx and subsequent brain clearance to the dcLNs. However, this observation requires further investigation to distinguish driving factors of this effect. Overall, K/Dex anesthesia promoted greater tracer accumulation within the dorsal skull, possibly reflecting isoflurane-induced brain volume changes (8-10) that may reduce dural tracer accumulation.

**Figure 3.**
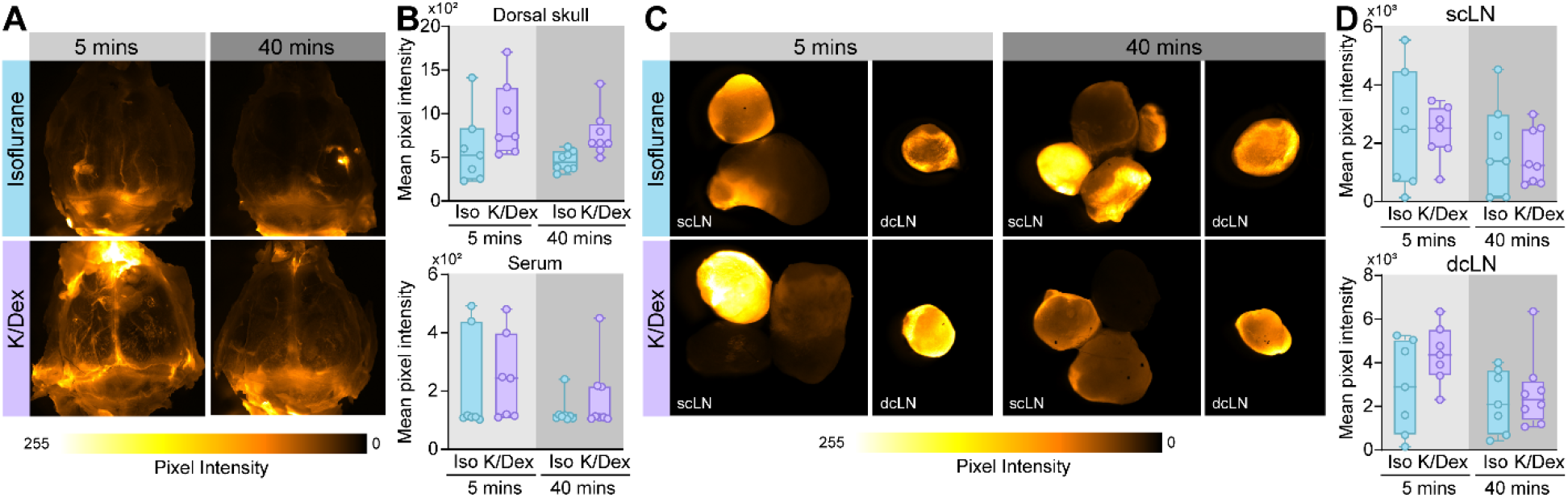
CSF efflux to dorsal skull, blood and cervical lymph nodes under isoflurane or ketamine/dexmedetomidine anesthesia. (A) Representative dorsal skull images obtained from mice under isoflurane or ketamine/dexmedetomidine (K/Dex) after a 5 minute (5 mins) or 40 minute (40 mins) intracisternal tracer infusion.(B) min/max boxplots showing mean fluorescent intensity in the dorsal skull (*top*) and blood serum (*bottom*). (C) Representative images of superficial cervical lymph nodes (scLN) and deep cervical lymph nodes (dcLN) under the same experimental conditions. Both left and right nodes were included in the quantification, although only one side is shown as representative images. (D) min/max boxplots showing fluorescent intensity in scLN (*top*) or dcLN (*bottom*). For all boxplots: the lower and upper whiskers indicate minimum and maximum values; the center line represent the median and individual circles represent individual mice. Statistical analysis was performed using two-way ANOVA, with exception of blood serum measurements which were analyzed using a Kruskal Wallis test. n = 7-8 mice per group. Complete statistical results are provided in the accompanying table.

## Discussion

This study demonstrates that isoflurane suppresses glymphatic influx compared with dexmedetomidine/ketamine independently of whether a 5- or 40-minute CSF tracer injection protocol is used (Fig. 1, 2). Efflux to the periphery, including cervical lymph nodes, was comparable across the 2 anesthetic regimens and 2 injection protocol (Fig. 3). These findings therefore support the large body of literature using the two anesthetic regimens as surrogates for wakefulness versus sleep (4-7), and argue against the notion that the reduced influx observed during isoflurane anesthesia is an artifact caused by washout of CSF tracers from the basal cisterns.

Studying brain fluid flow, and in particular the differences that exist across natural brain states, is inherently technically challenging because of the small body and brain size of mice and the need for invasive tracer administration. Even minor perturbations of physiological integrity can profoundly affect brain fluid dynamics; for example, glymphatic flow is suppressed for up to 24 hours after insertion of a small glass pipette into the brain (11). Similarly, the choice of anesthesia is critical, as anesthetics such as isoflurane or avertin suppress glymphatic flow, whereas α2-adrenergic agonist–based regimens, including dexmedetomidine- or xylazine-ketamine anesthesia, enhance CSF tracer influx (5).

Much of the current understanding of brain fluid transport, as well as the methodological refinements used to study it, has emerged progressively as the field has evolved and has since been validated across multiple laboratories (12). Nevertheless, the field remains relatively young, and its technical limitations must continue to be carefully acknowledged and experimentally addressed (13). In the present study, we found that the reduced CSF tracer influx observed during isoflurane anesthesia was not explained by rapid washout of tracers from the basal cisterns. Extending the tracer infusion duration from 5- to 40- minutes failed to eliminate the marked differences in glymphatic tracer influx between isoflurane and ketamine/dexmedetomidine anesthesia (Fig. 1, 2).

Isoflurane is a volatile halogenated general anesthetic that produces unconsciousness, immobility, amnesia, and muscle relaxation by broadly suppressing neuronal excitability throughout the central nervous system (14-16). Its actions are mediated through multiple molecular targets rather than a single receptor system. The best-established mechanism is potentiation of inhibitory neurotransmission through the GABA_A_ receptor. Isoflurane acts as a positive allosteric modulator of GABA_A_ receptors, enhancing chloride influx and thereby hyperpolarizing neurons and suppressing neuronal firing. At the same time, isoflurane suppresses excitatory neurotransmission. It inhibits NMDA-type glutamate receptors and reduces glutamatergic synaptic transmission, thereby decreasing cortical and thalamocortical excitability. Isoflurane additionally inhibits presynaptic neurotransmitter release through effects on voltage-gated sodium and calcium channels, reducing synaptic communication across neuronal networks. Another target class of isoflurane is the two-pore-domain potassium (K2P) channels, including TREK and TASK channels. Activation of these potassium channels hyperpolarizes neurons and further reduces neuronal excitability(17, 18).

At the systems level, isoflurane produces a characteristic brain state with reduced cortical activity, impaired thalamocortical communication, vasodilation, and altered autonomic regulation. Unlike α2-adrenergic agonist–based anesthesia, isoflurane suppresses slow-wave oscillatory activity and norepinephrine-dependent vascular dynamics that support glymphatic inflow (6, 19, 20). This mechanistic distinction likely contributes to the marked suppression of glymphatic CSF tracer influx observed during isoflurane anesthesia.

Ketamine/dexmedetomidine (K/Dex) anesthesia combines two mechanistically distinct agents that together produce sedation, analgesia, and a brain state that resembles several key features of natural non-rapid eye movement (NREM) sleep, including high-amplitude slow-wave EEG activity and reduced noradrenergic tone, both associated with glymphatic transport.

Dexmedetomidine is a highly selective α2-adrenergic receptor agonist that primarily acts on presynaptic autoreceptors in the locus coeruleus (LC), the major source of cortical norepinephrine (21). Activation of α2A receptors inhibits LC firing and suppresses norepinephrine release throughout the brain. This reduction in noradrenergic tone promotes synchronized slow-wave activity, decreases arousal, and induces a sleep-like sedative state that closely resembles endogenous NREM sleep (22). Because norepinephrine strongly regulates vascular tone, astrocytic volume, and extracellular space dynamics, suppression of norepinephrine signaling is thought to be a major mechanism by which dexmedetomidine enhances glymphatic influx (23). Reduced norepinephrine signaling is associated with expansion of the extracellular space, increased slow vasomotion, and enhanced CSF–interstitial fluid exchange (2, 23, 24).

Ketamine acts primarily as a noncompetitive NMDA receptor antagonist (25). By suppressing glutamatergic excitatory neurotransmission, ketamine contributes to anesthesia and analgesia while also reducing cortical activation. At subanesthetic or combined anesthetic doses, ketamine alone often produces dissociative and desynchronized cortical activity; however, when combined with dexmedetomidine, the strong α2-mediated suppression of LC activity dominates and generates stable slow-wave EEG oscillations (6). Ketamine additionally preserves cardiovascular function better than many anesthetics by maintaining sympathetic tone and arterial pulsatility to help sustain perivascular fluid transport.

At the systems level, K/Dex anesthesia therefore differs fundamentally from isoflurane anesthesia. Isoflurane broadly suppresses neuronal activity while inducing cerebral vasodilation and disrupting coordinated slow oscillatory vascular dynamics. In contrast, dexmedetomidine-based anesthesia recreates several hallmarks of natural sleep: low norepinephrine tone, synchronized slow oscillations, preserved vascular pulsatility, and enhanced glymphatic CSF influx (7, 26).

The findings reported here raise the broader question of why glymphatic flow is so strongly regulated by brain state. Although there is currently no direct evidence explaining why wakefulness suppresses glymphatic transport, we speculate that active large-scale fluid movement may not be compatible with the optimal operation of neural circuits in the awake brain. Fluid flow could increase glutamate spillover and thereby reduce the spatial and temporal fidelity of synaptic transmission required for precise sensory processing and rapid behavioral responses, rendering glymphatic flow in an awake animal potentially detrimental.

Nevertheless, multiple physiological processes that emerge during sleep appear to converge to support glymphatic transport. The sleeping brain is characterized by synchronized neuronal activity and high-amplitude slow-wave oscillations (2), and recent optogenetic studies have further shown that synchronization of neuronal activity enhances both CSF tracer influx and efflux (27). Similarly, fluctuations in cerebral blood volume are substantially larger during sleep than during wakefulness (28, 29). Because dynamic vascular volume changes are major drivers of glymphatic flow (23, 30), these enhanced sleep-associated vascular oscillations are likely to promote fluid transport through perivascular pathways.

In addition, the extracellular space expands during sleep (2, 24), likely in part as a consequence of reduced astrocytic cell volume, thereby lowering resistance to fluid movement through the brain parenchyma. Collectively, these and likely additional mechanisms either drive or facilitate the marked increase in glymphatic flow observed during sleep compared with wakefulness. Similarly, the distinct mechanistic actions of ketamine/dexmedetomidine versus isoflurane facilitate and suppress glymphatic flow respectively, independent of the CSF tracer injection paradigm.

## Methods

### Animals

Male and female wild-type, C57BL/6 mice (2-4 months old, 25-30 g) were used in all experiments. These mice were bred in the University of Rochester vivarium, with breeders refreshed each generation from Charles River Laboratories (Wilmington, MA). Animals were group-housed in a 12:12 light/dark cycle with ad libitum access to food and water. Animal holding rooms were maintained within temperature (70-74 degrees Fahrenheit) and humidity ranges (30–70%) ranges recommended by the ILAR Guide for the Care and Use of Laboratory Animals (1996). All efforts were made to minimize animal use and suffering. All experimental procedures were approved by the University of Rochester Medical Center Committee on Animal Resources.

### Drugs

Anesthesia for glymphatic influx experiments consisted of either 1.5-2.5% isoflurane in 100% O_2_, or 100 mg/kg racemic ketamine (Hikma, Berkeley Heights, NJ) and 1mg/kg dexmedetomidine (Tocris Biosciences, Bristol, U.K.) i.p. Depth of anesthesia was monitored using the pedal withdrawal reflex. If a response to toe pinch was observed, an additional one-tenth of the initial anesthetic dose was administered, and the tracer experiment was delayed until stable surgical anesthesia was achieved. Immediately prior to cisterna magna (CM) infusion, animal received one-tenth of the initial dose. Pedal reflexes were reassessed every 10□min throughout the experiments to ensure maintenance of adequate anesthetic depth during tracer circulation.

### Intracisternal CSF tracer infusion

Intracisternal tracer infusion were performed as described previously (5, 31, 32). Fluorescent CSF tracer consisted of Alexa Fluor ^™^ 647-conjugated bovine serum albumin, (66□kDa; Invitrogen, Life Technologies, Eugene, OR) was dissolved in artificial CSF at a concentration of 0.5% weight by volume (w/v). Anesthetized mice were secured in a stereotaxic frame, the CM surgically exposed, and a 30-gauge needle connected to PE10 tubing pre-filled with tracer was inserted into the CM. A total volume of 10 microliters of tracer was infused using a syringe pump (Harvard Apparatus, Holliston, MA) either at 2 microliters per minute for 5□minutes (5 mins injection) or 0.25 microliters per minute for 40 minutes (40 mins injection). Animals were euthanized by decapitation 40 min after initiation of tracer infusion, and tissues including brain, skull, blood, lymph nodes were collected. The brain was fixed overnight in 4% paraformaldehyde in PBS (Sigma-Aldrich, Burlington, MA).

Coronal vibratome slices (100□μm) were cut and mounted for imaging as previously described (33). Tracer influx was assessed ex vivo using conventional fluorescence microscopy of whole brains and coronal sections (Olympus, Stereo Investigator Software, Japan). Whole brain images, blood serum, and lymph node images were taken using a M205FA Leica fluorescent stereo microscope, coronal brain slices were imaged using Olympus MVX10 microscope.

Tracer influx was quantified by a blinded investigator using ImageJ software. For each section, the brain was manually outlined, and mean fluorescence intensity within the outline was measured. Regions of interest (ROIs) for dorsal skull analysis were restricted to areas surrounding the superior sagittal sinus (SSS) and transverse sinus. Average fluorescence intensity from each was used as a single biological replicate. Equivalent rostro-caudal levels were analyzed across all animals and no animals were excluded.

Representative images were selected from samples within one standard deviation of the group mean and processed identically across experimental groups to facilitate visualization of tracer distribution.

## Supporting information

Statistical Table

## Acknowledgments

The work was supported by National Institutes of Health (R01AT012312; R01AT011439, U19NS128613); Simons Foundation; Cure Alzheimer’s Fund, Ludwig Family Foundation; Sheldon G. Adelson Medical Research Foundation

